# Target sequence capture data shed light on the deeper evolutionary relationship on the subgenus Chamaecerasus of *Lonicera* (Caprifoliaceae)

**DOI:** 10.1101/2022.08.15.503957

**Authors:** Qing-Hui Sun, Diego F. Morales-Briones, Hong-Xin Wang, Jacob B. Landis, Jun Wen, Hua-Feng Wang

**Affiliations:** Sanya Nanfan Research Institute of Hainan University, Hainan Yazhou Bay Seed Laboratory, Sanya 572025, China; Department of Plant and Microbial Biology, College of Biological Sciences, University of Minnesota, 140 Gortner Laboratory, 1479 Gortner Avenue, Saint Paul, MN 55108, USA; Systematics, Biodiversity and Evolution of Plants, Department of Biology I, Ludwig-Maximilians-Universität München, Menzinger Str. 67, 80638, Munich, Germany; Zhai Mingguo Academician Work Station, Sanya University, Sanya 572022, China; School of Integrative Plant Science, Section of Plant Biology and the L.H. Bailey Hortorium, Cornell University, Ithaca, NY 14850, USA; BTI Computational Biology Center, Boyce Thompson Institute, Ithaca, NY 14853, USA; Department of Botany, National Museum of Natural History, MRC-166, Smithsonian Institution, PO Box 37012, Washington, DC 20013-7012, USA; Key Laboratory of Tropical Biological Resources of Ministry of Education, College of Tropical Crops, Hainan University, Haikou 570228, China

**Keywords:** *Lonicera*, Subgenus Chamaecerasus, Section Nintooa, Coeloxylosteum, Isika, Phylogenetics

## Abstract

The genus *Lonicera* L. is widely distributed and is well-known for its high species richness and morphological diversity. Previous studies have suggested that many sections of *Lonicera* are not monophyletic and phylogenetic relationships within the genus are still poorly known. In this study, we sampled 37 accessions of *Lonicera*, covering four sections of subgenus *Chamaecerasus* plus six outgroup taxa to recover the main clades of *Lonicera* based on sequences of nuclear loci generated by target enrichment and cpDNA from genome skimming. We found extensive cytonuclear discordance across the subgenus. Both nuclear and plastid phylogenetic analyses supported subgenus *Chamaecerasus* sister to subgenus *Lonicera*. Within subgenus *Chamaecerasus*, sections Isika and Niatoon were polyphyletic. Based on the nuclear and chloroplast phylogenies we propose to merge *Lonicera korolkowii* into section Coeloxylosteum and *Lonicera caerulea* into section Nintooa. In addition, *Lonicera* is estimated to have originated in the late Miocene (19.84 Ma). The stem age of section Nintooa was estimated to be 17.97 Ma (95% HPD: 13.31- 22.89). The stem age of subgenus *Lonicera* was estimated to be 16.35 Ma (95% HPD: 9.33- 45.15). Ancestral area reconstruction analyses indicate that *Lonicera* originated in the Qinghai Tibet Plateau (QTP) and Asia, with subsequent dispersal into other areas. The aridification of the Asian interior possibly promoted the rapid radiation of *Lonicera* within this region, and the uplift of the QTP appears to have triggered the dispersal and recent rapid diversification of the genus in the QTP and adjacent regions. Overall, this study provides new insights into the taxonomically complex lineages of *Lonicera* at the section level and the process of speciation.

## 1 Introduction

The discontinuous distribution, underlying formation mechanisms, and spatio-temporal evolution of plants in the north temperate zone have long been the focus of biogeographic research (Xiang et al., 201; Zhang et al., 2022). The boreotropical origin hypothesis suggests the presence of large forests in the middle and low latitudes of the Northern Hemisphere from the late Cretaceous to the early Eocene (Axelrod, 1973, 1975). With the decrease of global temperature, these forests gradually disappeared to form current distribution patterns (Lian et al., 2020; Zhang et al., 2022).

The land bridge hypothesis holds that the Bering Land Bridge and the North Atlantic Land Bridge provided a channel for the migration of species in Eurasia and North America. With the disappearance of the land bridges, species once widely distributed across Eurasia and North America formed the present discontinuous distribution patterns (Tiffney & Manchester, 2001). This has been supported by studies of *Cladrastis* Raf. (Fabaceae, Duan et al., 2020) and *Nyssa* Gronov. ex L. (Nyssaceae, Zhou et al., 2021). The ‘out-of-Tibet’ hypothesis holds that cold tolerant species dispersed from Tibet to the central and eastern part of China, Japan and North America after the climate became cold in the late Miocene, which has been supported in studies of Podophylloideae (Ye et al., 2022) and *Polygonatum* Mill. (Asparagaceae, Xia et al., 2022).

Currently, most studies on the discontinuous distribution of plants in the North temperate zone focus on the East Asia-North America discontinuous distribution (e.g., Wen, 1999; Donoghue et al., 2001; Zhou et al., 2021). However, little attention has been paid to the biogeographic study of the discontinuous distribution groups of Europe-Asia-North America. The plant genus Lonicera L. as is a typical representative of discontinuous distribution in the north temperate zone, providing an ideal model system for an in-depth study on the formation mechanism of the discontinuous distribution of plants in Europe-Asia-North America (Wen & Ickert-Bond, 2009).

*Lonicera* contains about 180 species distributed in North Africa, Asia, Europe and North America, of which 57 can be found in China (Yang et al., 2011). *Lonicera* is an evergreen or deciduous upright shrub or dwarf shrub with single leaves, opposite, sparse whorls, and no stipules. The flowers are frequently paired with axillary, 5-lobed, neat or lip-shaped calyx and corolla, and form a berry (Fig.1). Honeysuckle plants are not only medicinally important, but also are drought-tolerant and excellent soil and water conservation or dust-retaining plants. The majority of the fruits are edible and have a high ornamental value. Some skin types are a good source of fiber. Plant fibers can be turned into artificial cotton, flowers can be extracted into fragrant oils, and seed oil can be turned into soap (Hsu & Wang, 1988; Mabberley, 2008).

**Figure 1.**
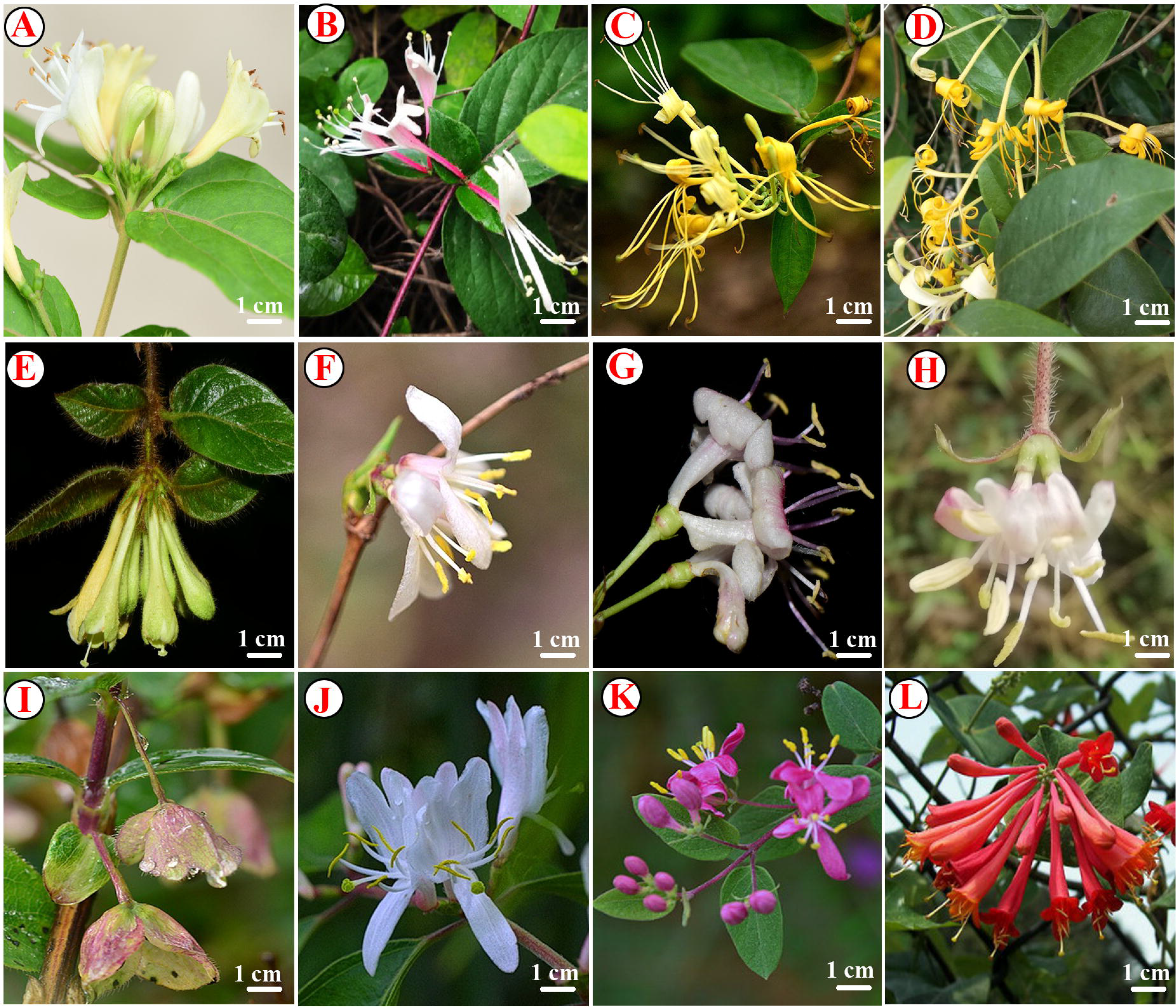
Photographs of *Lonicera* taxa. (A) *L.macrantha* (D.Don) Spreng (Nintooa); (B) *L. japonica* Thunb.(Nintooa); (C) *L. hypoglauca* Miq.(Nintooa); (D) *L. confusa* DC.(Nintooa); (E) *L. reticulata* Champ.(Nintooa); (F) *L. fragrantissima* Lindl. & J. Paxton(Isika); (G) *L. gynochlamydea* Hemsl.(Isika); (H) *L. fragrantissima* var. *lancifolia* Lindl. & J. Paxton(Isika); (I) *L. hispida* Pall. ex Schult. (Isika); (J) *L. maackii* (Rupr.) Maxim.(Coeloxylosteum); (K) *L. tatarica* L. (Coeloxylosteum); (L) *L. rupicola* Hook. f. & Thomson (Isoxylosteum).

Since Linnaeus (1753) established *Lonicera*, the genus has historically posed various systematic problems, especially with infrageneric classification. Maximowicz (1877) was the first to systematically classify Asian *Lonicera* and then subdivide the genus into three subgenera [*Caprifolium* (Tourn.) Maxim., *Chamaecerasus* (Tourn.) Maxim. and *Xylostron* (Tourn.)]. Subsequently, Rehder (1903, 1913) further divided the subgenus *Chamaecerasus* into four sections and 20 subsections. Nakai (1938) classified Japanese *Lonicera* species into 15 sections and 8 subsections within the genus *Lonicera*. Hara (1983) classified *Lonicera* into two subgenera [Lonicera and Caprifolium (Mill.) Dippel], f our sections [Isika (Anderson) Rehder, Caeruleae (Rehder) Nakai, Lonicera and Nintooa (Sweet) Maxim] within the subgenus *Chamaecerasus*, and five subsections within section Isika. Hsu and Wang (1988) proposed a new system for Chinese *Lonicera* species by splitting the genus into two subgenera. They also identified four sections and 12 subsections within the subgenus *Chamaecerasus* in China. Conversely, Yang et al. (2011) did not recognize taxa at the section level in their Flora of China treatment.

Nakaji (2015) used morphological data and various DNA markers to explore the phylogenetic relationships within *Lonicera*, such as chloroplast DNA (cpDNA) *rpo*B-*trn*C, *atp*B-*rbc*L, *trn*S-*trn*G, *pet*N-*psb*M, *psb*M-*trn*D; and nuclear ribosomal DNA (nrDNA) ITS. Theis (2008) used sequences of the ITS region of nuclear ribosomal DNA and five chloroplast non-coding regions (*rpo*B-*trn*C spacer, *atp*B-*rbc*L spacer, *trn*S-*trn*G spacer, *pet*N-*psb*M spacer, and *psb*M*trn*D spacer) to address these problems. However, until now, the circumscription of *Lonicera* and its phylogenetic position have long been problematic and controversial. Despite previous phylogenetic studies, the relationships among sections, subsections and species of *Lonicera* (Caprifoliaceae), especially in China, remain unclear. In this context, we explored the phylogenetic relationships and biogeographical diversification of *Lonicera* subgenus *Chamaecerasus* using a targeted enrichment datasets of nuclear loci and plastomes from genome skimming.

## 2 Materials and methods

### 2.1 Sampling, genomic data generation

We sampled 32 species from subgenus *Chamaecerasus*, four species from subgenus *Lonicera*, and six families that were included as outgroups [*Symphoricarpos orbiculatus* Moench, *Heptacodium miconioides* Rehd., *Weigela florida* (Bunge) A. DC., *Diervilla lonicera* Mill., *Sambucus williamsii* Hance, and *Viburnum opulus* var. *americanum* Aiton]. Species, vouchers and GenBank accession numbers are given in Table 1.

**Table. 1.**
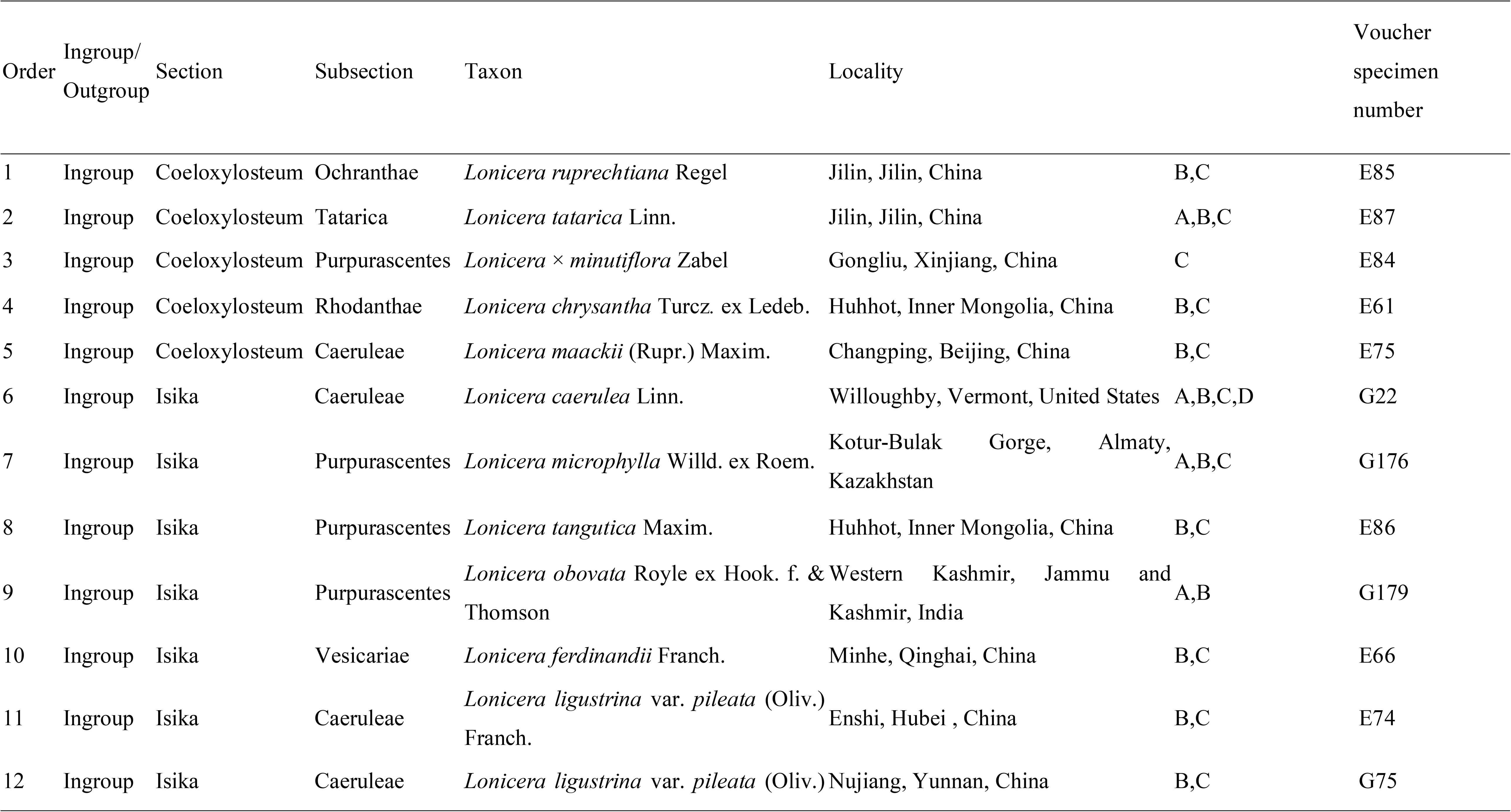
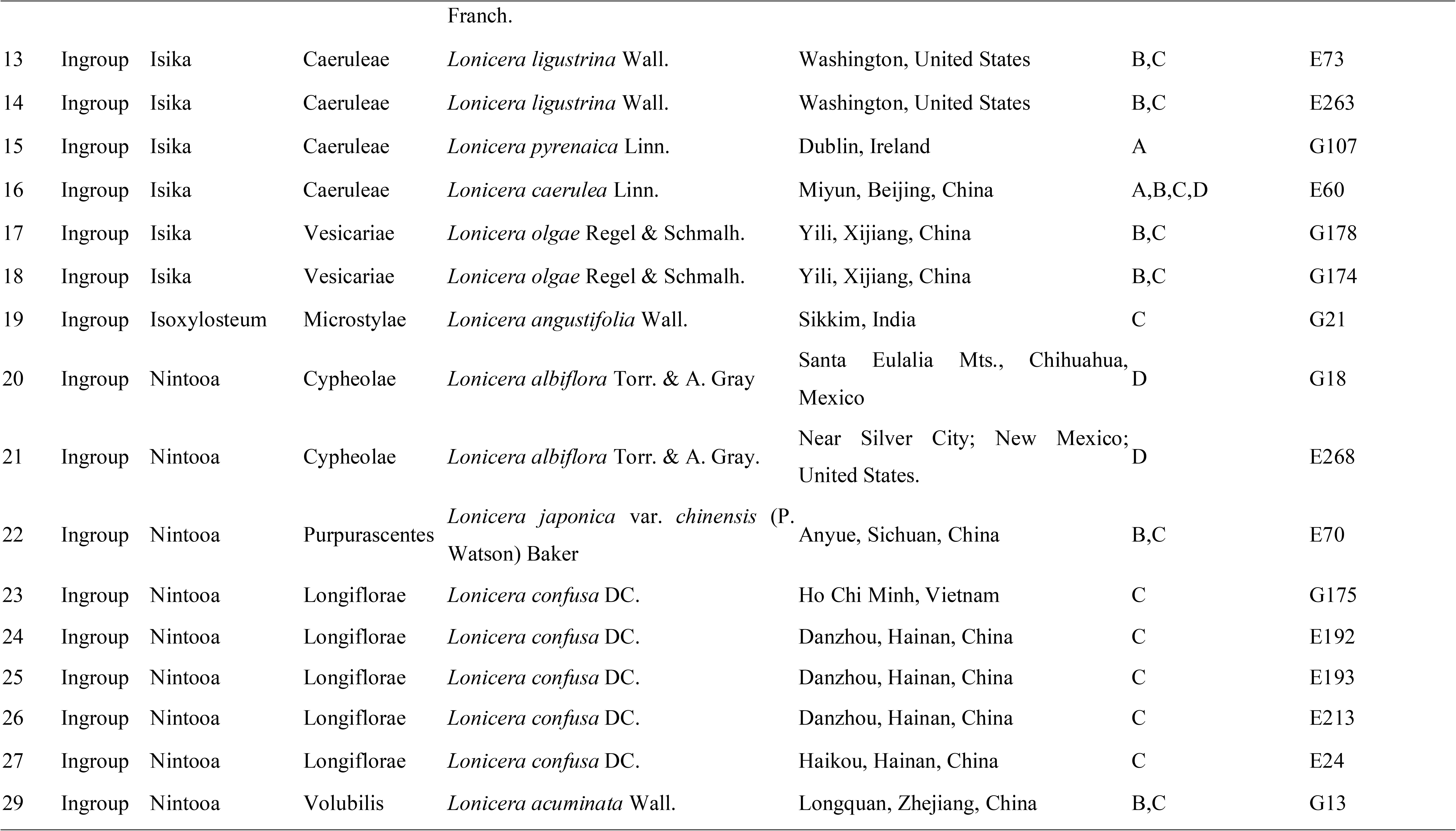
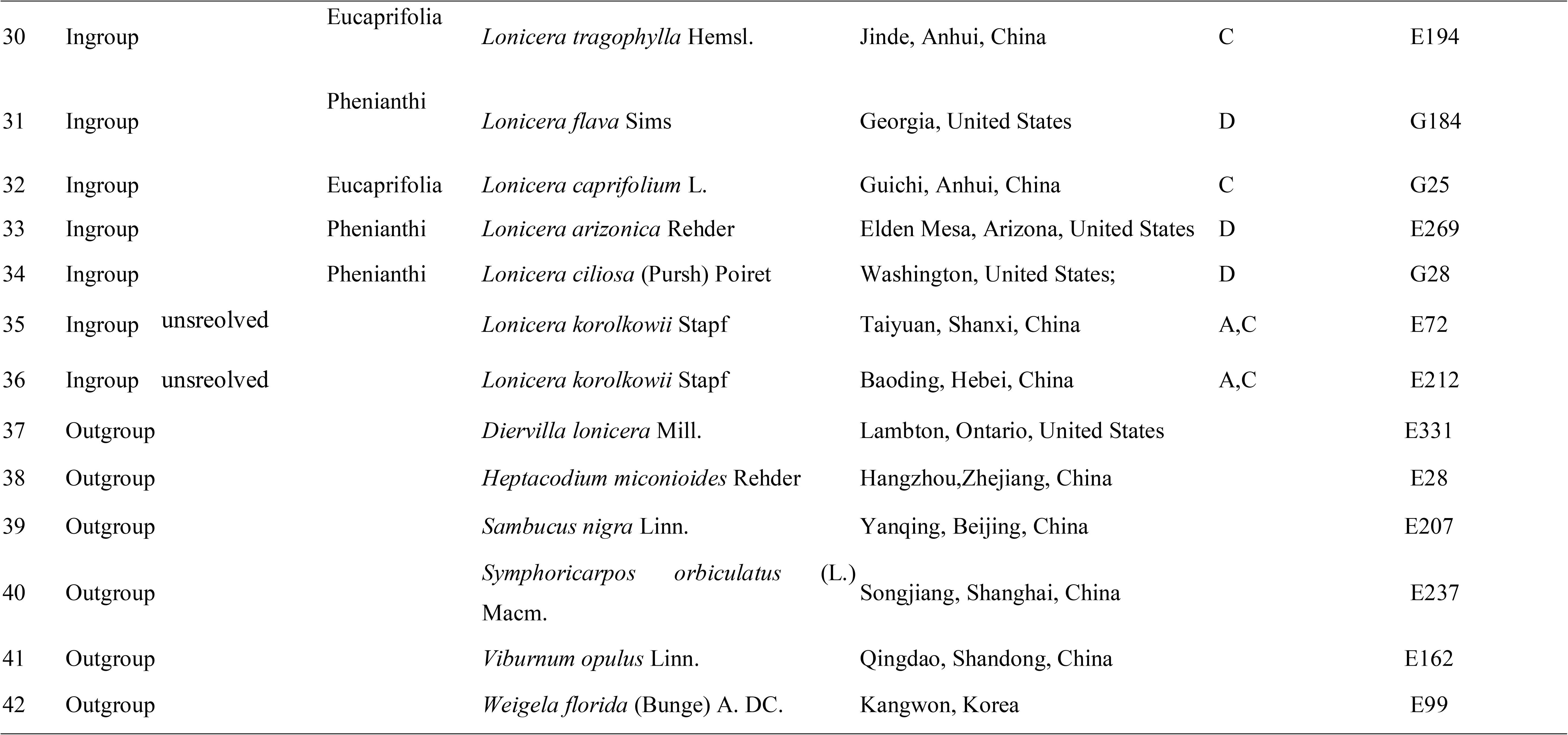
List of species and vouchers used in this study.

We extracted DNA from dried leaf tissue using a modified cetyltrimethylammonium bromide (CTAB) method (Doyle and Doyle, 1987). We used a target enrichment approach (Weitemier et al., 2014) to sequence 428 putatively single-copy nuclear genes using baits design across Dipsacales (Wang et al., 2021). Hybridization, enrichment, and sequencing methods followed those of Wang et al. (2021). We obtained full plastomes from genome skimming data with detailed descriptions of plastome sequencing methods in Wang et al. (2020a, b).

### 2.2 Read processing and assembly

Raw reads were cleaned using Trimmomatic v.0.36 (Bolger et al., 2014) by removing adaptor sequences and low-quality bases (ILLUMINACLIP: TruSeq ADAPTER: 2:30:10 SLIDINGWINDOW: 4:5 LEADING: 5 TRAILING: 5 MINLEN: 25). Nuclear loci were assembled with HybPiper v.1.3.1 (Johnson et al., 2016). Exon assemblies were carried out individually to avoid chimeric sequences in multi-exon genes produced by potential paralogy (Morales-Briones et al., 2018) and only exons with a reference length of more than 150 bp were assembled. Paralog detection was performed for all exons with the ‘paralog investigator’ warning. To obtain ‘monophyletic outgroup’(MO) orthologs (Yang and Smith, 2014), all assembled loci (with and without paralogs detected) were processed following Morales-Briones et al (2020).

For plastome assembly, raw reads were trimmed using SOAPfilter v.2.2 (BGI, Shenzhen, China). Adapter sequences and low-quality reads with Q-value ≤ 20 were removed. Clean reads were assembled against the plastome of *Kolkwitzia amabilis* (KT966716.1) using MITObim v.1.8 (Hahn et al., 2013). Assembled plastomes were preliminarily annotated with Geneious R11.04 (Biomatters Ltd., Auckland, New Zealand) to detect possible inversions/rearrangements, with corrections for start/stop codons based on published *Lonicera* plastomes. Detailed descriptions of plastome assembly methods can be found in Wang et al. (2020a).

### 2.3 Phylogenetic analyses

#### 2.3.1 Nuclear dataset

We analyzed the nuclear data set using concatenation and coalescent-based methods. We used IQ-TREE v. 1.6.1 (Nguyen et al., 2015) for Maximum likelihood (ML) analysis (Table 2). We used Quartet Sampling (QS; Pease et al., 2018) to distinguish strong conflict from weakly supported branches. We used ASTRAL-III v.5.7.1 (Zhang et al., 2018) for species tree estimation, with the input being individual exon trees generated with RAxMLv8.2.12 **(Stamatakis, 2014)** using a GTR+G. Detailed descriptions of phylogenetic analyses methods can be found in Wang et al. (2020a).

**Table 2.**
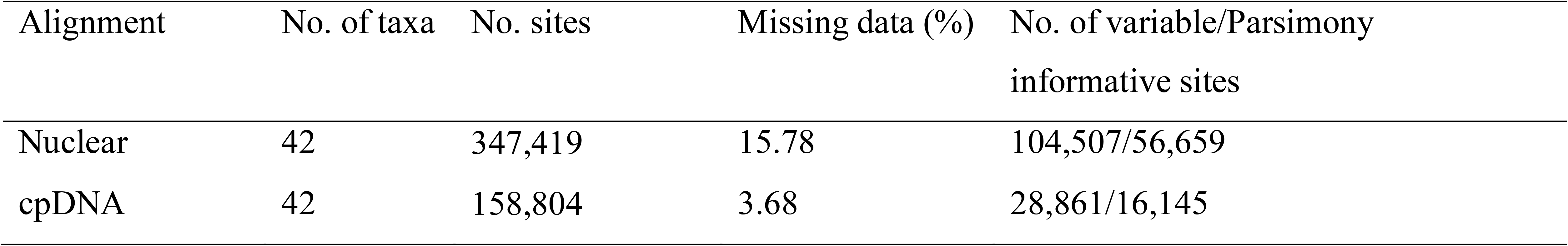
Dataset statistics, including the number of taxa, number of characters, number of PI characters, missing data.

#### 2.3.2 Plastome dataset

Complete plastomes were aligned with MAFFT v.7.407 (Katoh and Standley, 2013) and aligned columns with more than 90% missing data were removed using Phyutility (Smith and Dunn, 2008). We performed extended model selection (Kalyaanamoorthy et al., 2017) followed by ML gene tree inference and 200 non-parametric bootstrap replicates for branch support in IQ-TREE v.1.6.1 (Nguyen et al., 2015).

### 2.4 Species network analysis

We reduced the 42 samples data set to one outgroup and 14 ingroup taxa representing all major clades based on the nuclear analyses. We looked for evidence of hybridization (reticulation) within *Lonicera* via species network search carried out with the InferNetwork MPL function in PhyloNet (Wen et al., 2018). We used the command ‘CalGTProb’ in PHYLONET to calculate the probability scores of the concatenated RAxML, ASTRAL, and cpDNA trees, given the individual gene trees, to estimate the optimal number of hybridizations and measure if the species network matched our gene trees better than a purely bifurcating species tree. Finally, we selected the best model using the bias-corrected Akaike information criterion (AICc; Sugiura, 1978), Akaike information criterion (AIC; Akaike, 1973), and the Bayesian information criterion (BIC; Schwarz, 1978). The network with the lowest AICc, AIC and BIC scores were selected as the best-fit-species network.

### 2.5 Divergence time estimation

The nuclear dataset was used to estimate the divergence time of *Lonicera* with BEAST v.2.4.0 (Bouckaert et al., 2014). We employed the GTR+G substitution model and a Birth-Death model for tree priors. Other parameters were left at default values. We selected the available fossils as calibration points, the fossil seeds of *Weigela* Thunb. from the Miocene and Pliocene in Poland (Lańcucka-Rodoniowa, 1967), and the Miocene in Denmark (Friis, 1985) to constrain the stem age (offset 23.0 Ma, lognormal prior distribution 23.0-28.4 Ma). The root age was set to 78.9 Ma (mean 78.9 Ma, normal prior distribution 76.3-82.2 Ma) following Li et al. (2019). We performed multiple MCMC runs for a total of 50 billion generations sampling every 5000 generations. Stationarity was checked with Tracer v.1.7 (Rambaut et al., 2018). All ESS values exceeded 200, and the first 25% of trees were discarded as burn-in. We performed the burn-in and combined trees from independent runs in LogCombiner, and then determined the maximum clade credibility (MCC) tree with mean heights for the nodes in TreeAnnotator The 95% highest posterior density (HPD) intervals were viewed in FigTree v.1.4.2 (Drummond et al., 2012).

### 2.6 Ancestral area reconstructions

The MCC tree obtained from the nuclear data was used for biogeographical analysis in RASP v.3.21 (Yu et al., 2015). We used the dispersal-vicariance-analysis (DIVA) model (Ree & Smith, 2008). Four major areas of distribution were categorized: (A) Europe; (B) Qinghai Tibet Plateau; (C) Asia; and (D) North America. Each sample was assigned to its respective area according to its contemporary distribution range (Table 1). The BBM analyses ran for 100 million generations using 10 MCMC chains.

### 2.7. Ecological niche modelling

We used MAXENT v.3.2.1 (Phillips et al., 2006) to perform Ecological Niche Modelling (ENM) for *Lonicera*. We used 19 bioclimatic variables (Table S1) as environmental data for the ENM analysis. The data included baseline climate (BioClim layers for the period 1950 −2000 at a spatial resolution of 30 s arc), data for the last glacial maximum (LGM; ~22,000 years BP) with spatial resolution at 2.5 min arc resolution simulated by Community Climate System Model (CCSM), Max Planck Institute Model (MPI) and Model for Interdisciplinary Research on Climate (MIROC), and the last interglacial period. Twenty-five replicate runs were performed in MAXENT to ensure reliable results. The MAXENT jackknife analysis was used to evaluate the relative importance of the 19 bioclimatic variables employed. All ENM predictions were visualized in ArcGIS v.10.2 (ESRI, Redlands, CA, USA).

### 2.8. Analysis of character evolution

To evaluate the evolution of discrete characters, we used the RAxML trees (based on nuclear and cpDNA data) with the character mapping options in Mesquite v.3.51 (Maddison and Maddison 2018). The Markov k-state one-parameter model of evolution for discrete unordered characters (Lewis, 2001) was used. We selected two morphological characters to reconstruct the evolution of morphological features. The character states were encoded according to Alfred (1903) and Landrein (2017). Fruit color were scored with the following: (0) Red or (1) Blue black. Seed coat was scored with the following: (0) Pilose, (1) Hairless, (2) Pubescent or (3) Bristle.

## 3. Results

### 3.1 Exon assembly

The number of assembled exons per sample ranged from 251 (*Lonicera caerulea* E60) to 923 (*Lonicera tangutica* E86) (with > 75% of the target length), with an average of 766 exons (Table S1; Fig. S1). The number of exons with paralog warnings ranged from five in *Lonicera caerulea* E60 to 275 in *Lonicera olgae* G178 (Table S1). The concatenated alignment had a length of 347,419 bp, including 56,659 parsimony-informative sites, and a matrix occupancy of 82% (Table 2). The cpDNA alignment resulted in a matrix of 158,804 bp with 16,145 parsimony-informative sites and a matrix occupancy of 95% (Table 2).

### 3.2. Phylogenetic reconstruction

We recovered the *Lonicera* phylogeny based on nuclear and cpDNA data sets (Figs. 2 and 3). For the nuclear concatenated tree, cpDNA tree and Astral tree (Fig. 2 and 3), subgenus *Chamaecerasus* is sister to subgenus *Lonicera*. Our nuclear phylogenetic analyses recovered four sections Isika, Nintooa, Coeloxylosteum, and Isoxylosteum of subgenus *Chamaecerasus* with maximum support (BS = 100). Based on the nuclear concatenated tree (Fig. 2, left), we found section Coeloxylosteum forming a clade with *Lonicera korolkowii* (BS=100), and section Coeloxylosteum (including *Lonicera korolkowii*) was sister to sections Isoxylosteum + some members of Isika. Nintooa is not monophyletic in the cpDNA tree since Isoxylosteum is embedded in the section. Furthermore, we see that sections Isika and Nintooa are polyphyletic. Section Nintooa forms a clade with *Lonicera caerulea* (BS=100). Subgenus *Lonicera* forms a clade with *Lonicera albiflora* (BS=89), which is then sister with subgenus *Chamaecerasus*.

**Figure 2.**
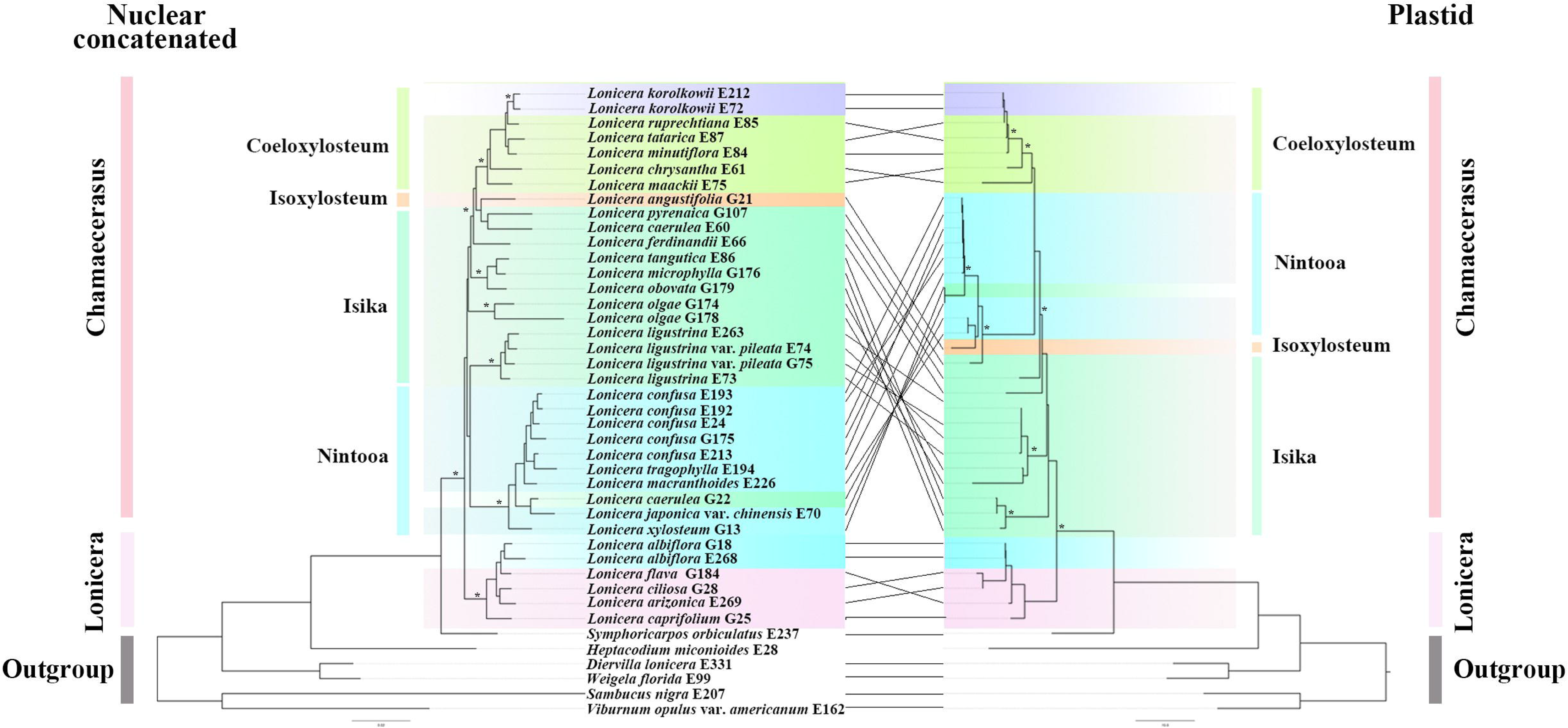
Tanglegram of the nuclear concatenated (left) and plastid (right) phylogenies. Dotted lines connect taxa between the two phylogenies. Support values are shown above branches. The asterisks indicate maximum likelihood bootstrap (BS) support of 100%. Major taxonomic groups or main clades in the family as currently recognized are indicated by branch colors as a visual reference to relationships.

**Figure 3.**
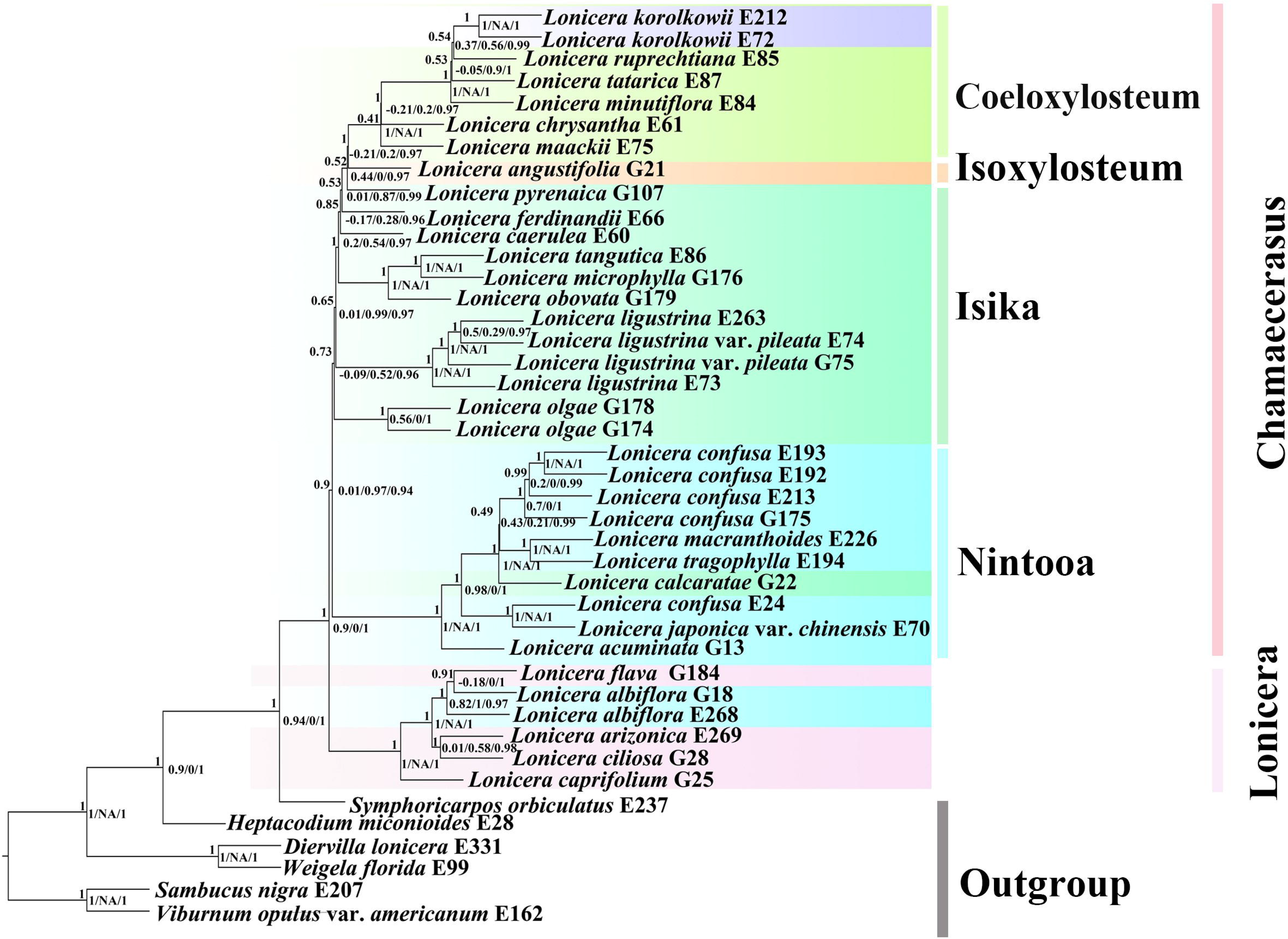
ASTRAL-III species tree. Clade support is depicted as: Quartet Concordance (QC)/Quartet Differential (QD)/Quarte Informativeness (QI)/Local posterior probabilities (LPP).

For the cpDNA tree (Fig. 2, right), we found section Coeloxylosteum forms a clade with *Lonicera korolkowii* (BS=100), and section Coeloxylosteum (including *Lonicera korolkowii*) is sister to some members of section Nintooa + section Isoxylosteum + members of section Isika. Section Nintooa forms a clade with *Lonicera caerulea* (BS=99), and section Nintooa (including *Lonicera caerulea*) was Isoxylosteum is embedded within the section, along with one species of Isika. Section Isika was polyphyletic. Subgenus *Lonicera* forms a clade with *Lonicera albiflora* (BS=90), which is in turn sister with subgenus *Chamaecerasus*.

For the Astral tree (Fig. 3), we found subgenus *Lonicera* forms a clade with *Lonicera albiflora* (LPP=1), and in turn is sister with subgenus *Chamaecerasus* with strong QS support but a signal of a possible alternative topology (0.9/0/1). Section Coeloxylosteum forms a clade with *Lonicera korolkowii* (LPP=0.54), and section Coeloxylosteum (include *Lonicera korolkowii*) is sister to sections Isoxylosteum + Isika with QS counter support as a clear signal for alternative topology (−0.21/0.2/0.97). Section Nintooa forms a clade with *Lonicera caerulea* (LPP*=*0.9), and is in turn sister with sections Coeloxylosteum + Isoxylosteum + Isika with moderate QS support (0.01/0.97/0.94).

In our species network analyses, we inferred three reticulation events as the best networks in the 15 samples tree (Table3, Figs.4, S2). Within this best network model, the first reticulation event involved the clade with *Lonicera angustifolia* G21, the inheritance probabilities analysis showed that the major genetic contributor came from the clade of *Lonicera tatarica* E87 and *Lonicera minutiflora* E84 (58.4%). The second reticulation event involves the clade of *Lonicera confusa* and *Lonicera caerulea*, with the major genetic contributor (92.7%) coming from their ancestor. The third reticulation event involves the clade of *Lonicera tatarica* E87, *Lonicera minutiflora* E84, *Lonicera pyrenaica* G107, *Lonicera tangutica* E86, *Lonicera microphylla* G176. The major genetic contributor (72.0%) came from the clade of *Lonicera ciliosa* G28 and *Lonicera albiflora* E268 (Fig.4).

**Figure 4.**
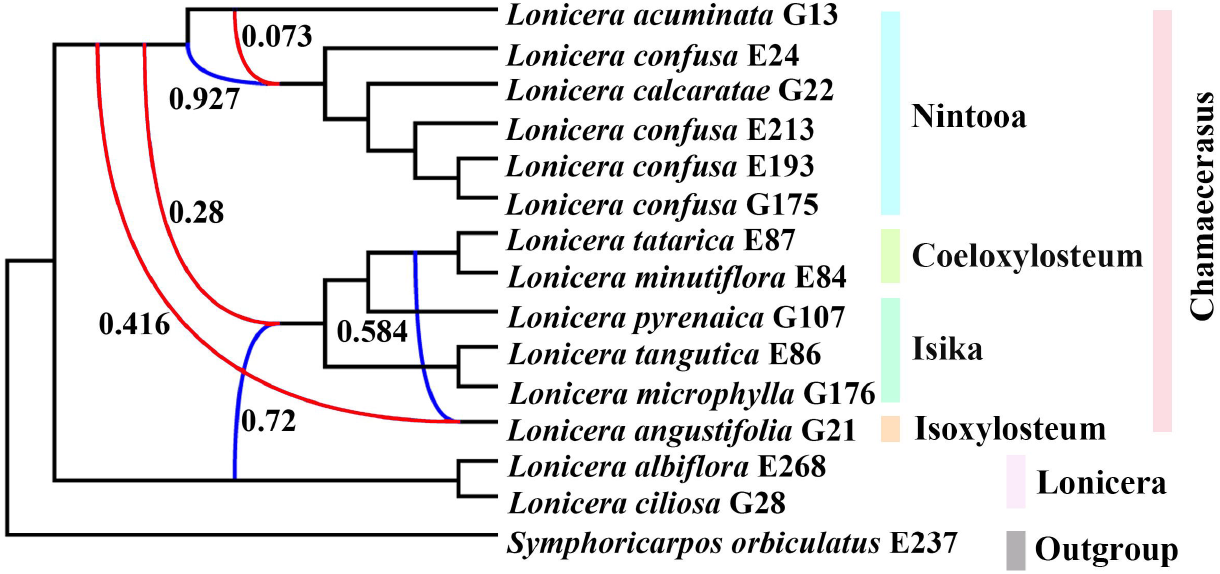
Best supported species networks inferred with PhyloNet for the 15 samples. Numbers next to the inferred hybrid branches indicate inheritance probabilities. Red lines represent minor hybrid edges (edges with an inheritance contribution < 0.50).

**Table 3.**
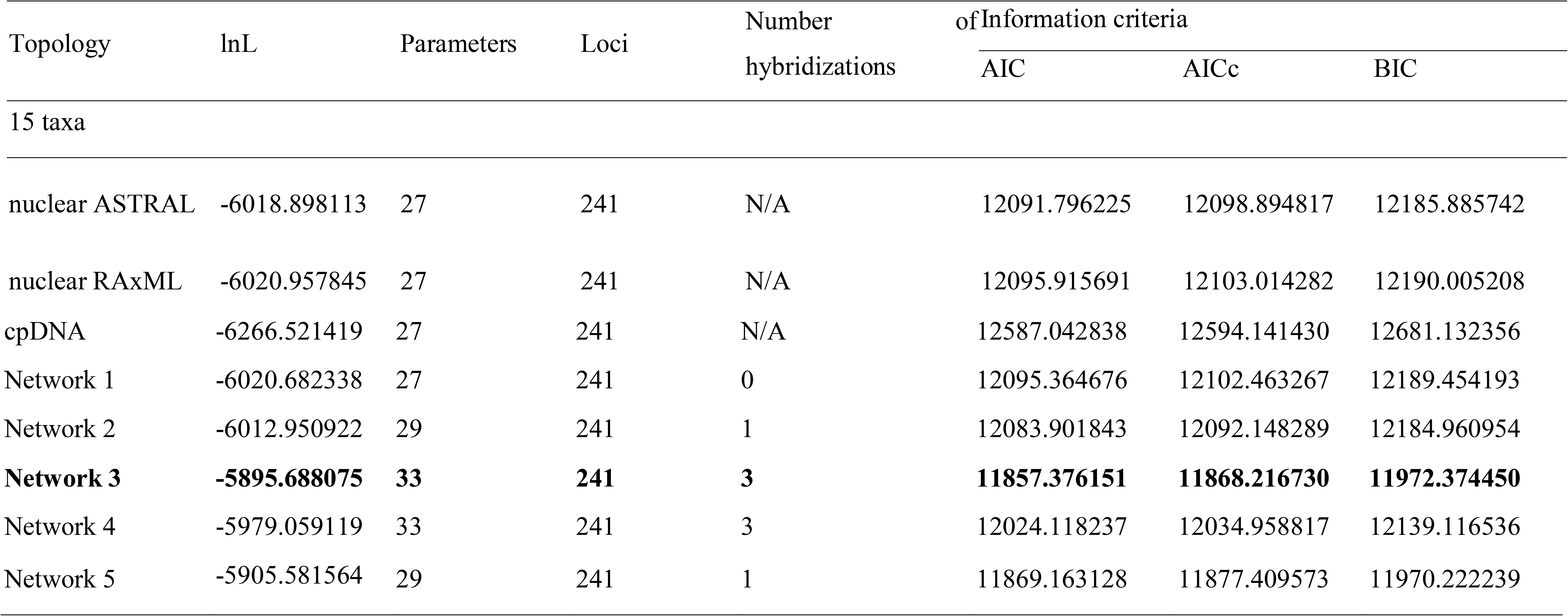
Model selection of the different species networks and bifurcating trees.

### 3.2 Divergence time estimation

Divergence time estimates for the genus *Lonicera* using the nuclear data are presented in Fig.5. The extant genus *Lonicera* dated to the late Miocene (19.84 Ma, node 2) (95% HPD: 13.74-61.89; node 2). The stem age of section Nintooa was estimated to be 17.97 Ma (95% HPD: 13.31-22.89; node 3). The split age of subgenus *Lonicera* was estimated to be 16.35 Ma (95% HPD: 9.33-45.15; node 5). The stem age of section Coeloxylosteum was estimated to be 13.65 Ma (95% HPD: 10.32-18.22; node 7).

**Figure 5.**
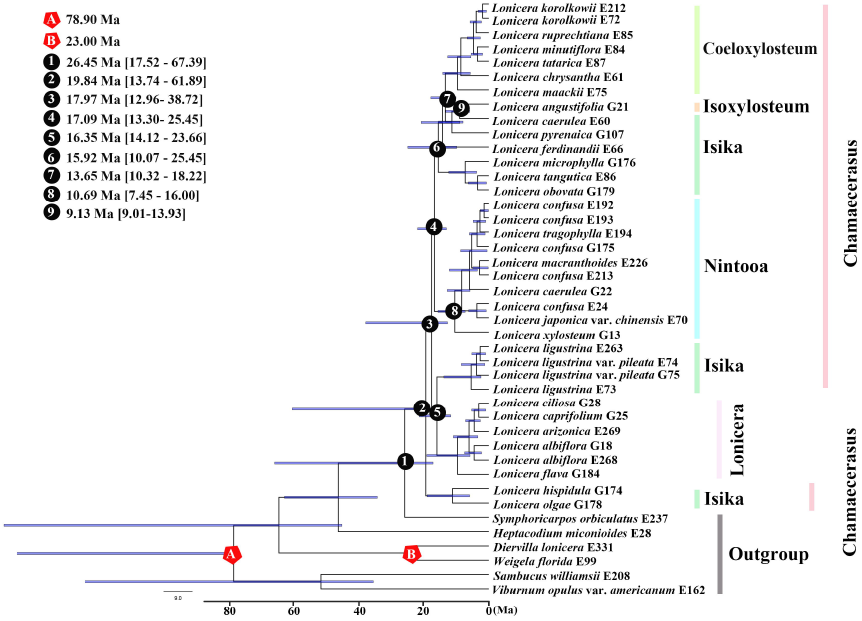
BEAST analysis of divergence times based on nuclear data. Calibration points are indicated by stars. Numbers 1-9 represent major divergence events in *Lonicera*. Mean divergence times and 95% highest posterior densities are provided for each major clade.

### 3.3 Ancestral area reconstruction

According to our ancestral area reconstructions paired with divergence time estimation, genus *Lonicera* originated around the late Miocene in Asia (Fig. 6). Our analyses revealed 19 dispersal events and three vicariance events among the four defined biogeographic areas (Fig. 6). The ancestor of *Lonicera* was inferred to have been present throughout the Qinghai Tibet Plateau (region B) and Asia (region C), with subsequent dispersal or vicariance across other areas [Europe (region A) and North America (region D)].

**Figure 6.**
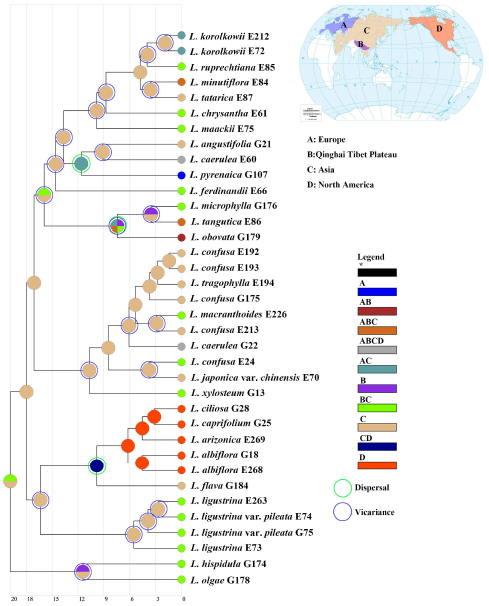
Ancestral area reconstruction for *Lonicera*. Areas of endemism are as follows: (A) Europe; (B) Tibet (Qinghai Tibet Plateau); (C) Asia; (D) North America.

### 3.3 Ecological niche modelling and niche identity tests

The AUC test result (mean ± SD) for the ENM, averaged across all 25 runs, was significant (0.996 ±0.000 for section Isoxylosteum, 0.993± 0.001 for section Isika, 0.995± 0.001 for section Coeloxylosteum, 0.993± 0.001 for section Nintooa). In our analysis, the LGM climate and the present time climate species distribution areas show different distribution patterns. We used CCSM, MIROC and MPI models to simulate the LGM climate for the four sections of subgenus *Chamaecerasus*, and the inferred distribution from the two LGM models (CCSM and MPI models) were highly similar. For the CCSM and MIROC models (the MIROC model area simulation during the LGM period is warmer and drier than the CCSM model), the distribution density and range of subgenus *Chamaecerasus* under the MPI model was slightly lower than the those generated from the CCSM model, but the differences were not significant (Fig. 7 and S3). Compared with the LGM climate, the potential distribution range of subgenus *Chamaecerasus* species in Mainland China will be much smaller, while the habitat remains stable in most of the present time climate distributions.

**Figure 7.**
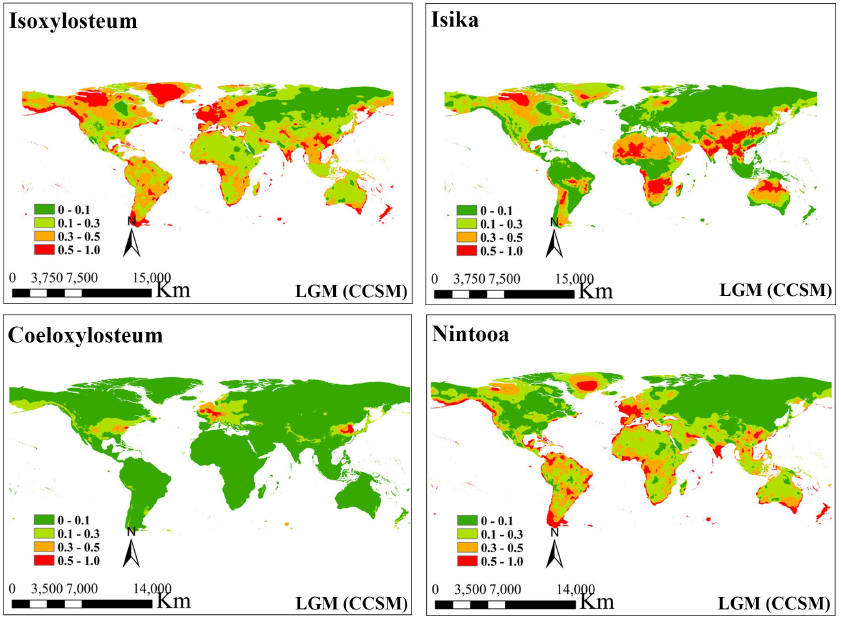
Comparison of potential distributions as probability of occurrence for sections Isoxylosteum, Isika, Coeloxylosteum and Nintooa at the CCSM climatic scenarios of the Last Glacial Maximum (LGM, ca. 21,000 years BP). The maximum training sensitivity plus specificity logistic threshold has been used to discriminate between suitable (cutline 0.1-1 areas) and unsuitable habitat. The darker color indicates a higher probability of occurrence.

### 3.4 Character evolution

Character mapping of two qualitative traits on the ML tree (using both nuclear and cpDNA) demonstrated that fruit characters are homoplasious within *Lonicera* (Figs. S4, S5). The ancestral fruit color for *Lonicera* is red for both the nuclear (Fig. S4) and cpDNA (Fig. S5). The nuclear tree shows two origins of blue-black fruit color, while the cpDNA tree shows a single origin of blue-black fruits. In the nuclear analysis, our results suggest that seed coat have little homoplasy, but the ancestral character of coat was most likely pubescent for the entire tree but hairless for *Lonicera* (Fig. S4). In the cpDNA analysis, our results suggest that seed coat has high homoplasy, the ancestral state for coat character for the entire tree was uncertain due to the diversity in types among the major early-diverging clades (Fig. S5). However, the ancestral state for *Lonicera* is still a hairless coat. Overall, we found that the patterns of character evolution from the nuclear gene tree and cpDNA tree were similar.

## 4. Discussion

### 4.1 Phylogenetic conflicts between cytonuclear genomes

Angiosperm phylogeny has been studied extensively using genome-scale data sets (Gitzendanner et al., 2018; Leebens-Mack et al., 2019; Li et al., 2019). The relationships between Lonicera, on the other hand, are still a mystery. It is clear from these examples that simply increasing taxon sampling will not lead to more accurate inference or increased bootstrap support for problematic deep nodes in the angiosperm phylogeny.

Any amount or quality of molecular data can’t explain why certain genes’ phylogenetic relationships are in constant discordance. Lineage sorting (ILS) that fails to fix shared ancestral variation between closely spaced speciation events is frequently to blame for this discrepancy. Conflicting phylogenetic signals can be caused by a group’s recent history and ancestral variation being sorted out in this way. [e.g., African rift cichlids (Brawand et al.,2014), Platyfish (Cui et al., 2013), and Horses (Jónsson et al.,2014)] and at deeper timescales [major land plant families (Wickett et al., 2014) and major bird lineages (Jarvis et al., 2014)]. A second emerging pattern is that post-speciation hybridization (introgression) appears to be substantially more commonplace than previously appreciated. A diverse range of animal groups all show evidence of post-speciation gene flow [e.g., Butterflies (Martin et al., 2013), Horses (Garrigan et al., 2014), Fish (Cui et al., 2013), Flies (JÓ nsson et al., 2014), Mosquitoes (Fontaine et al., 2015), and Galápagos finches (Lamichhaney et al., 2015)]. The frequency and extent of introgression is remarkable given that introgressive hybridization has played little role in conventional models of animal speciation and diversification.

ILS and post-speciation introgression contribute to generating more complex evolutionary histories than can be represented by simple bifurcating trees. In response, new approaches are being developed to account for these potential sources of gene tree discordance, including the use of MP-EST (Liu et al., 2010) and ASTRAL (Mirarab et al., 2014) to infer the underlying species. Diversity studies of beak size differentiation in Galápagos finches have successfully clarified ambiguous species relationships and highlighted loci that might underpin specific functional or ecological changes that accompanying rapid phylogenetic transitions (Lamichhaney et al., 2015).

We recovered the *Lonicera* phylogeny based on nuclear and cpDNA data sets (Figs. 2 and 3), our results indicate that subgenus *Lonicera* (including *Lonicera albiflora*) is sister to subgenus *Chamaecerasus*. Additionally, sections Coeloxyloseum (including *Lonicera korolkowii*) was monophyletic, while sections Nintooa (including *Lonicera caerulea*) and Ishika were polyphyletic, this result is consistent with previous studies (Theis et al., 2008; Nakaji et al., 2015). In this study we do not have the power to determine if section Isoxyloseum is monophyletic because only one sample was included. Thus, more data are needed to test the monophyly of section Isoxyloseum.

For the species network analyses, our results show that the parental contributions for the reticulation events detected in *Lonicera* (Fig. 4) are unequal. Within subgenus *Chamaecerasus*, section Nintooa clade, the inheritance contributions (0.073 and 0.927) support the existence of extensive gene flow in section Nintooa. For section Isoxylosteum (*L. angustifolia* G21), the inheritance contributions (0.416 and 0.584) support a hybridization event between the ancestral lineage of the section Coeloxylosteum clade and the ancestral lineage of the section Nintooa clade. For the ancestral lineage of the sections Coeloxylosteum + Isika clade, the inheritance contributions (0.28 and 0.72) support the introgression event between the ancestral lineage of the section Nintooa clade and the ancestral lineage of the subgenus *Lonicera* clade. Notably, all our network results show that all events overlap with the parent population section Nintooa, suggesting that there is extensive hybridization and infiltration in section Nintooa, which may be the main reason that the section Nintooa and subsection classified according to morphology do not form a monophyly.

### 4.2 Dating estimation and biogeographic analyses

Smith & Donoghue (2010) previously reported the divergence time of subgenus *Chamaecerasus* in the northern hemisphere to be 7-17 Ma. Temperature and precipitation were keys to limiting the survival of the ancestral plants of subgenus *Chamaecerasus* (Smith, 2008, 2009). In this study, the stem age of *Lonicera* was dated to the late Miocene, ca. 19.84 Ma (Fig. 5, node 2), later than the early uplift of the Qinghai-Tibet Plateau (QTP). This result is similar to Smith (2008), who found that *Lonicera* originated from Asia in the late Miocene or late Oligocene, and was widely distributed in Asia, Europe and western North America in the Oligocene.

Based on our ancestral area reconstructions (Figs. 6; Table 1), *Lonicera* represents a typical case of a species-rich plant genus that originated from the QTP and Asia and subsequently migrated into other regions. As such, *Lonicera* is consistent with the hypothesis of ‘out-of-Tibet’ (Mao et al., 2010; Jia et al., 2012; Zhang et al., 2012), suggesting that the common ancestor of *Lonicera* during the late Miocene was adapted to higher-altitude and cold environments. Dispersal and rapid diversification of the group was likely triggered by the global climatic cooling in the late Miocene (Mao et al., 2010). Due to the uplift of the QTP, the temperature in the interior of Asia decreased and humidity increased, making plant distributions in Asia appear to rapidly distribute (Budantsev, 1987, 1993; Chen et al., 2021; Ye et al., 2022). Global cooling provides a dominant driving factor for the drying of central Eurasia because it reduces the moisture load of the westerly winds and decreases precipitation in inland Asia (Lu et al., 2010; Miao et al., 2012). Some species scattered in Europe may pass through the mountains of Central Asia and Asia Minor (Jia et al., 2012; Jiang et al., 2019). As summarized by Wu et al. (2018), the rise of dryland flora in different continents almost began in the middle and late Miocene, indicating that this period is a key period for the aggregation and evolution of modern dryland flora. A total of 19 independent dispersal events of were detected, indicating extensive hybridization/introgression within *Lonicera*, possibly undergoing rapid radiation.

### 4.3 Morphological characters evolution

The characters of stamen number and fruit colour are traditionally used for generic recognition within Caprifoliaceae (Backlund, 1996; Donoghue et al., 2003; Yang & Landrein, 2011; Landrein et al., 2020). Due to variable environmental selective pressures or other phenomena, inconsistencies between morphological characteristics may occur. The evidence indicates that seed coat has little homoplasy, and the patterns of character evolution from nuclear gene tree and cpDNA tree were similar.

Members of section Nintooa are vines and their upper leaves are reduced or even bractlike. There are three subsections, including the monotypic subsection Calcaratae, which is unique in having a long nectar spur, connate ovaries, and bracteoles. The explanation for this unresolved phylogeny may be incomplete lineage sorting of ancestral polymorphisms before the rapid divergence of these species, making plastid DNA sequences insufficient to resolve the phylogenetic relationships. Section Isika is the largest section in *Lonicera*, and it is distributed throughout the entire range of the genus. This group has the most diverse character (e.g., habit, size and shape of the bracts and bracteoles, external scales of the winter buds and the shape of the corolla). Section Coeloxylosteum is a rather homogeneous group, characterized by evanescent pith, distinctly two-lipped corollas, and the tendency to have distinct ovaries, an upright habit, and red fruits. Two subsections, Ochranthae and Tataricae, are distinguished only by minor morphological differences. Section Isoxylosteum are compact shrubs with solid, white pith, small leaves, and flowers with nearly actinomorphic corollas that are non-gibbous at the base.

In the systems of Hara (1983) and Ohba (1993), section *Caeruleae* (=Isika), characterized by fused ovaries and connate bracteoles, is represented by *Lonicera caerulea* (= *L. coerulea). Lonicera caerulea* is a deciduous shrub. Inflorescences axillary, paired flowers; bracteoles glabrous, fused into a cupule tight-enclosing and free ovaries. Corolla tubular-funnelform, outside puberulent, shallowly gibbous at base. However, in our analysis, *Lonicera caerulea* is nested within section Niatoon, with a well-supported topology (Figs. 2 and 3).

In the system of Rehder (1903 and 1913), *Lonicera albiflora* is classified as section Niatoon. *Lonicera albiflora* is a deciduous shrub. The pollen is red, opposite to the axils of leaves, fragrant. Nowadays, we find that botanists have not classified *Lonicera korolkowii*. Our phylogenetic relationship shows that subgenus *Lonicera* forms a clade with *Lonicera albiflora*, with a well-supported topology (Figs. 2 and 3).

At present, all classification systems of *Lonicera* do not classify *Lonicera korolkowii. Lonicera korolkowii* is a deciduous shrub. The pollen is red, opposite to the axils of leaves, fragrant. Our phylogenetic relationships show that Section Coeloxylosteum forms a clade with *Lonicera korolkowii*, with strong supported topology (Figs. 2 and 3).

Researchers have proposed their own classification systems (Hsu and Wang, 1988; Yang and Landrein, 2011) based on the evolutionary law of different organs or characters of *Lonicera*. Due to the limited number of samples and experimental means, there were defects in each classification system. A solid phylogenetic framework is the premise and foundation for exploring the evolutionary history and speciation mechanism of species. Overall, this study recovered the phylogeny of *Lonicera* through the nuclear and cpDNA data sets, we obtained the generally well-supported topology.

### 4.4 Taxonomic treatment

*Lonicera albiflora*, Torrey & Gray, Fl. N. Am. 2: 6 (1841). - Gray, Jour. Nat. Hist. Boston 6: 213 (1857); Syn. Fl. N. Am. 12: 18 (1884). **Subgenus *Lonicera*.**

*Lonicera caerulea*, L., 1: 174. 1753. (1 May 1753) (Sp. Pl.). Switzerland: Habitat in Helvetia. **Section Nintooa**.

*Lonicera korolkowii*, Stapf, Gard. & Forest 7: 34 (1894). **Section Coeloxylosteum**.

## Conclusions

Natural hybridization/introgression is extensive in *Lonicera* and may require further investigation with extensive populational sampling. Our findings support the use of Hyb-Seq data at the section level to clarify the boundaries among four sections. For other taxa, our study demonstrates the importance to use both nuclear and plastid genome data, as well as network analysis, to untangle the potential reticulation history. Our results also indicate that genus *Lonicera* dated to the late Miocene (19.84 Ma). Additionally, our biogeographic analyses support the ‘out-of-Tibet’ hypothesis. This work better resolves the deep phylogenetic relationships among the main lineages within subgenus Chamaecerasus, particularly the relationships between the four currently recognized sections within subgenus Chamaecerasus.

## Author contributions

H.F.W. and J.W. conceived the study. H.F.W. and D.F.M-B. performed the research and analyzed the data. Q.H.S., D.F.M-B., H.X.W., J.W., J.B.L, and H.F.W. wrote and revised the manuscript.

## ACKNOWLEDGEMENTS

The work was funded by Hainan Provincial Natural Science Foundation of China (421RC486), the Project of Sanya Yazhou Bay Science and Technology City (Grant number: SCKJ-JYRC-2022-83), and National Science Foundation of China (31660055). We appreciate Gabriel Johnson for his help with the target enrichment experiment, and the United States National Herbarium for collection access. We acknowledge the staff in the Laboratories of Analytical Biology at the National Museum of Natural History, the Smithsonian Institution for technical support and assistance.

---

**Fig. S1.**
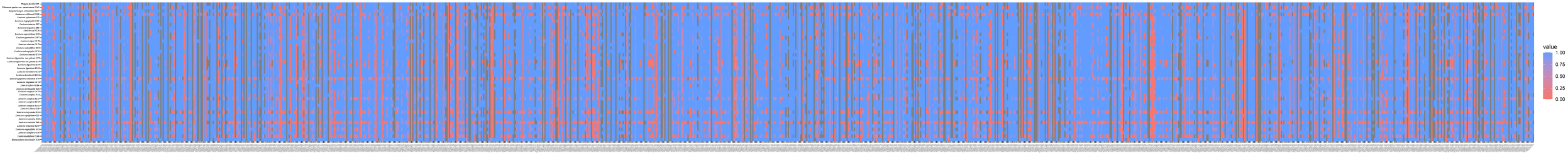
Heatmaps showing gene recovery efficiency for the nuclear gene in 43 species of *Lonicera*. Columns represent genes, and each row is one accession. Shading indicates the percentage of the reference locus length coverage.

**Fig. S2.**
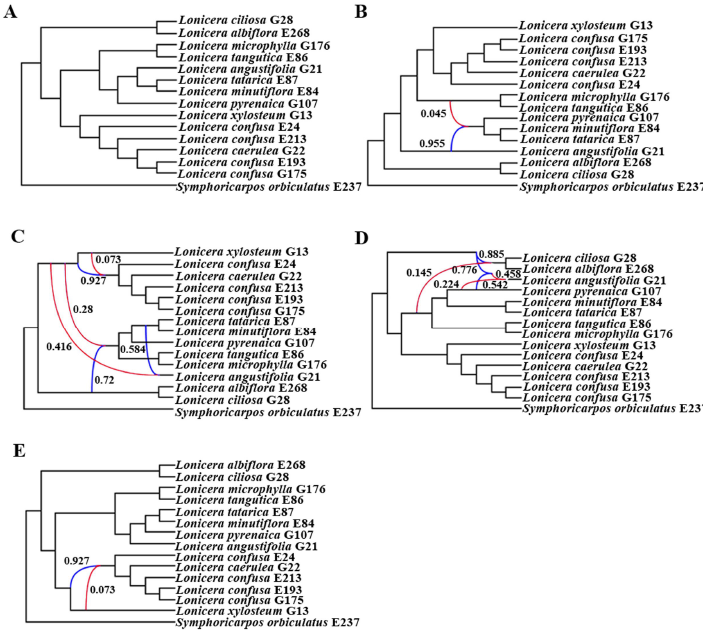
Best species networks of the selective nuclear data estimated with PhyloNet for the 15 samples data set. A: One hybridization event; B: Two hybridization events; C: Three hybridization events; D: Four hybridization events; E: Five hybridization events. Blue branches connect the hybrid nodes. Numbers next to blue branches indicate inheritance probabilities.

**Fig. S3.**
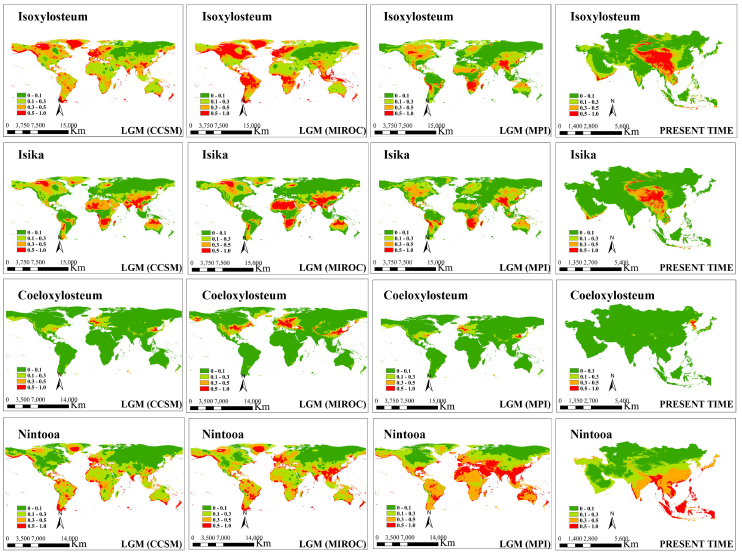
Potential distribution probability (in logistic value) of occurrence for *Lonicera* in the world generated form MAXENT.

**Fig. S4.**
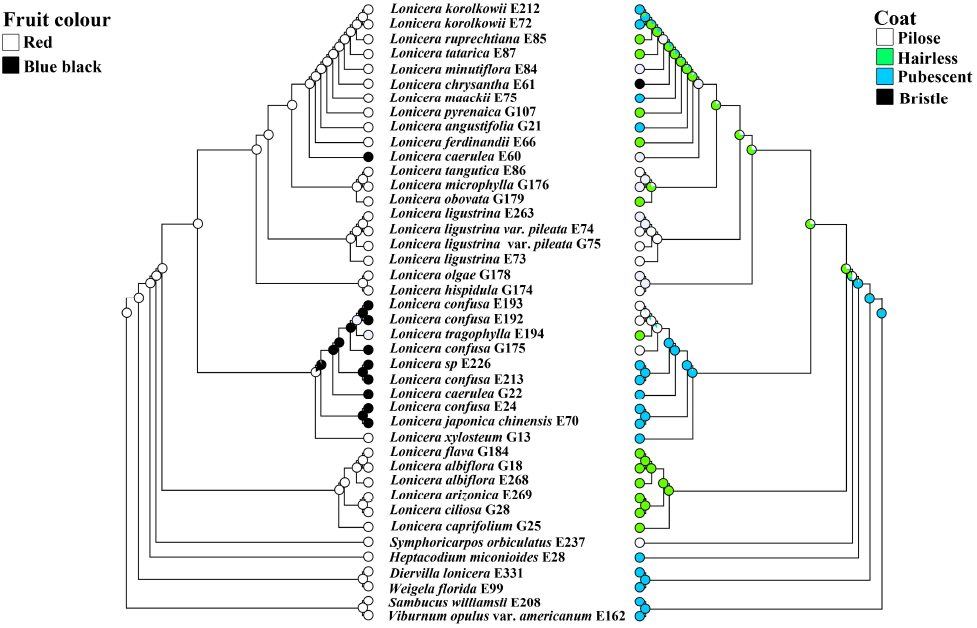
Likelihood inference of character evolution in *Lonicera* using Mesquite v.2.75 based on nuclear matrix. Left, Fruit color; Right, Coat.

**Fig. S5.**
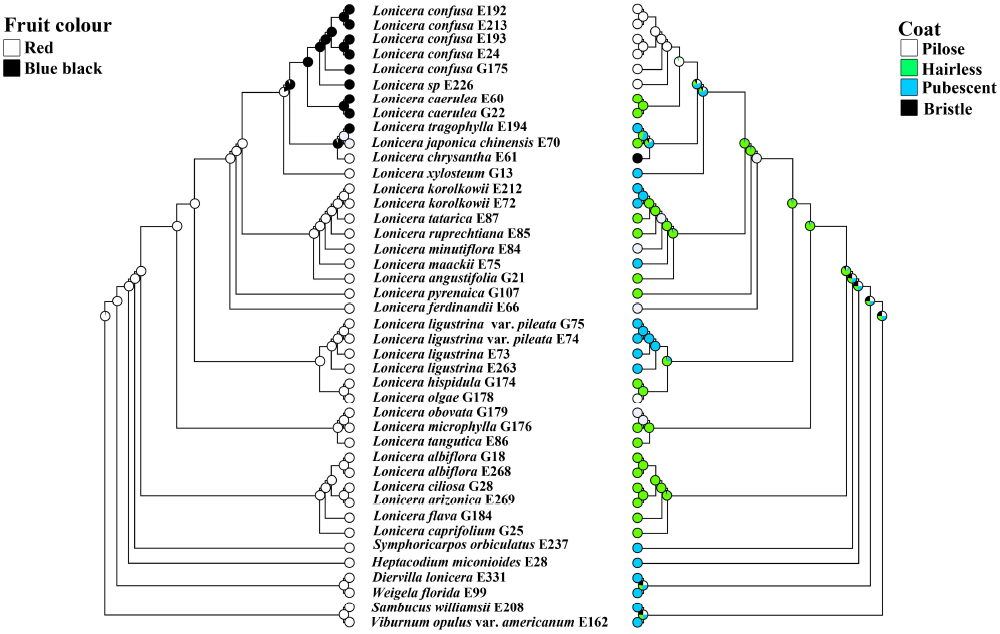
Likelihood inference of character evolution in *Lonicera* using Mesquite v.2.75 based on plastid matrix. Left, Fruit color; Right, Coat.

**Table. S1.**
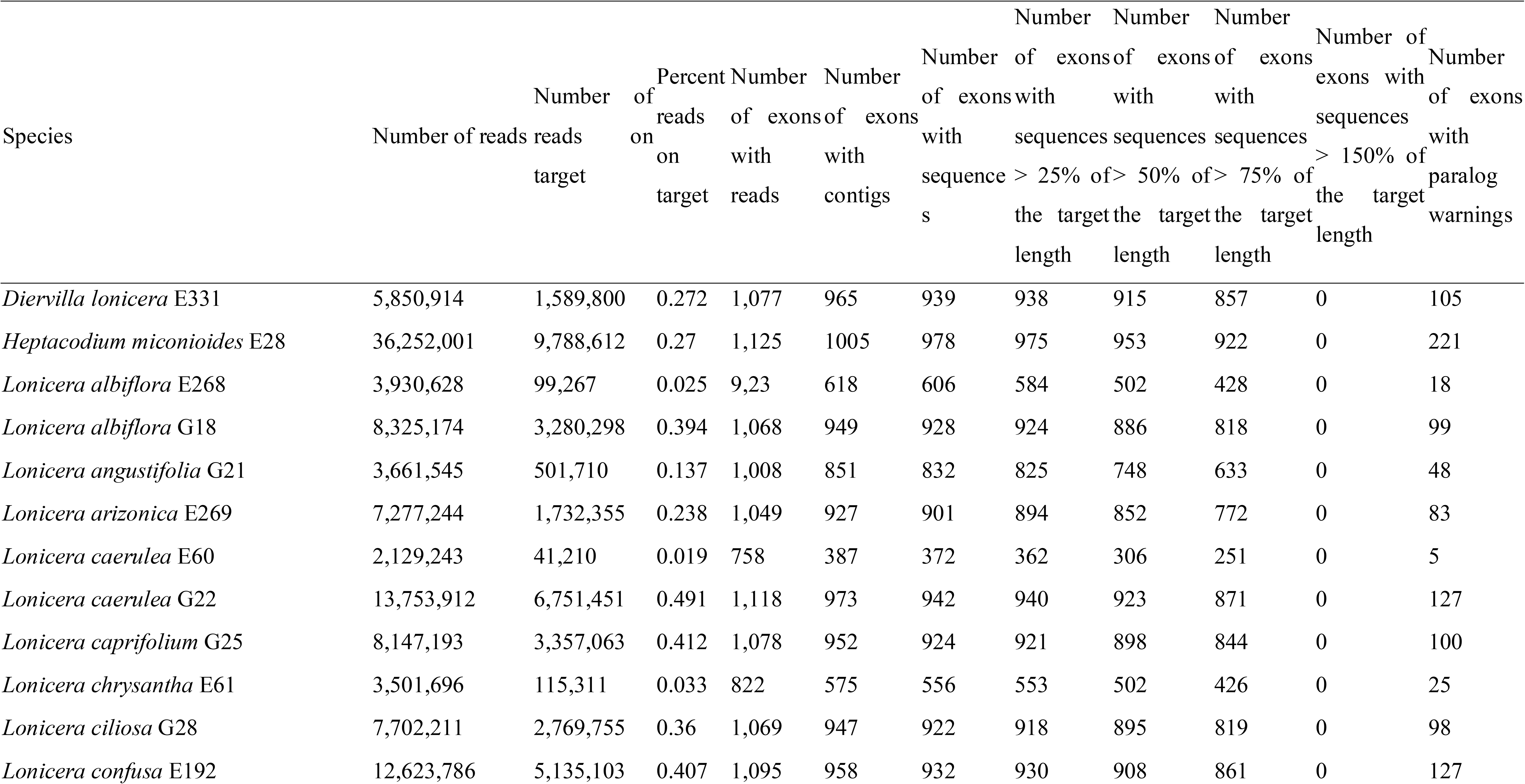
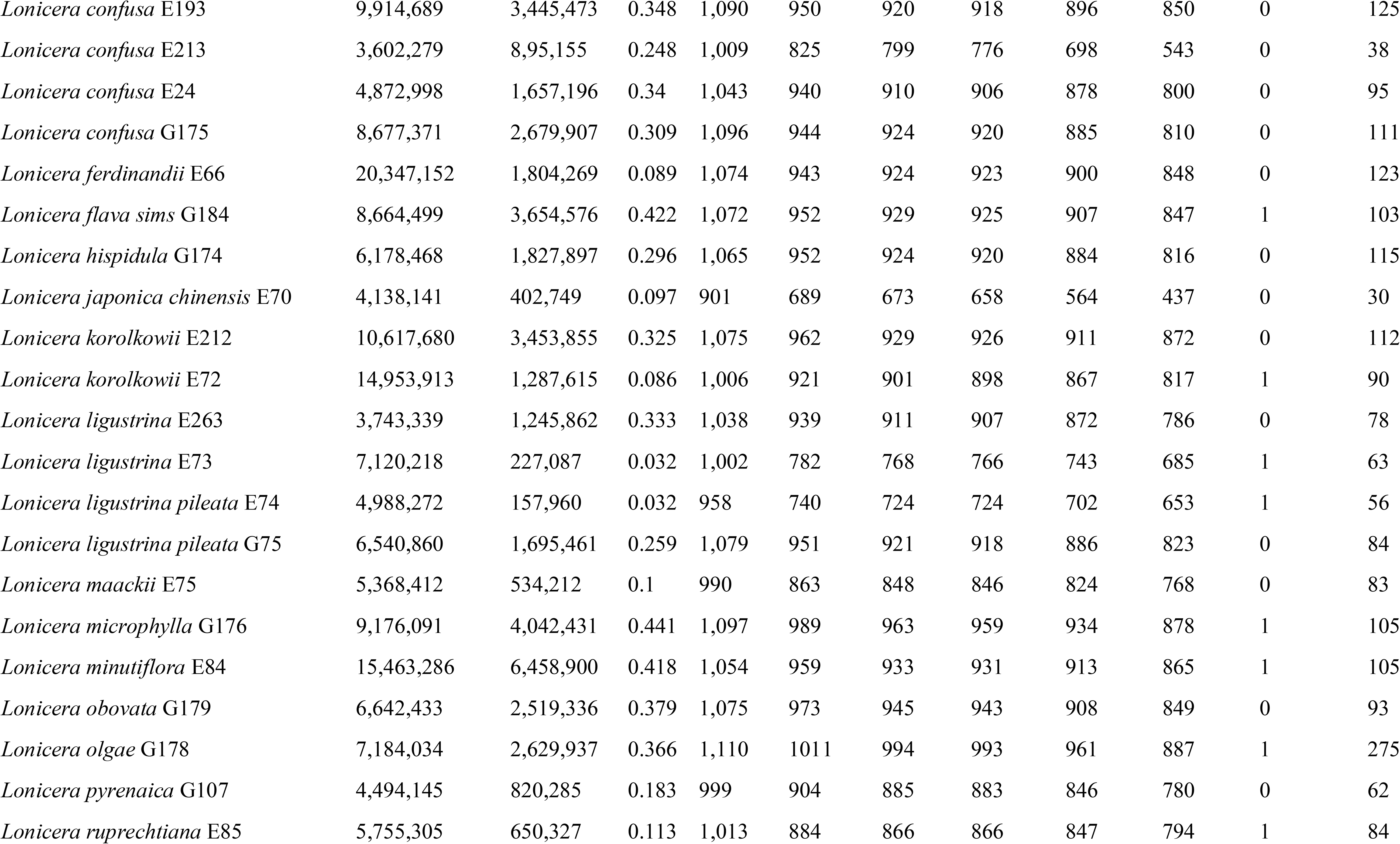
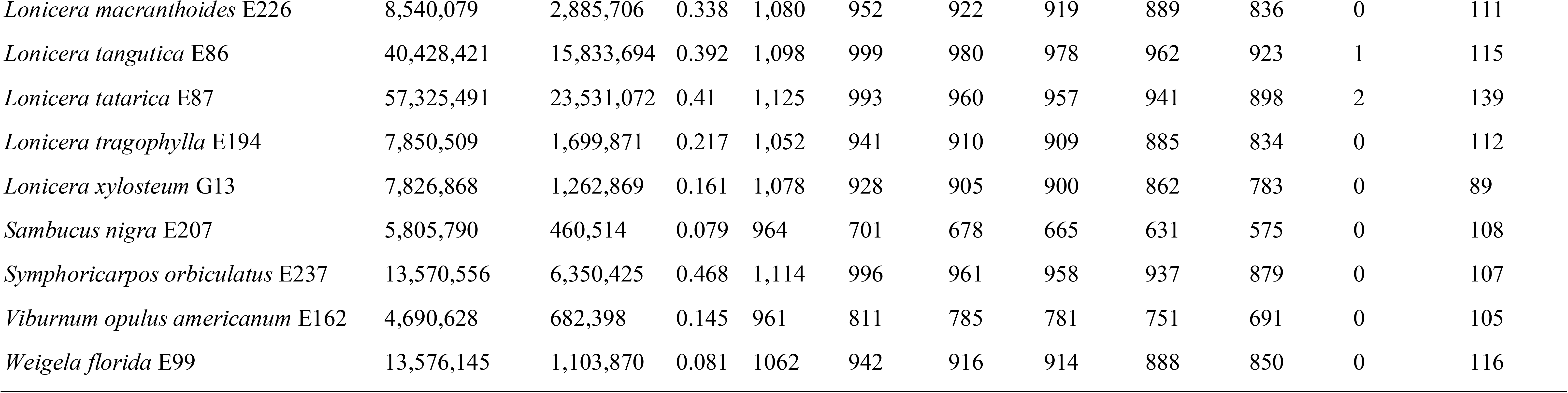
The 19 bioclimatic variables for ENM analyses.

Table. S2 HybPiper assembly statistics.

